# Hyperspectral characterisation of natural illumination in woodland and forest environments

**DOI:** 10.1101/2021.07.19.452949

**Authors:** Li Shiwen, Laura Steel, Cecilia A. L. Dahlsjö, Stuart N. Peirson, Alexander Shenkin, Takuma Morimoto, Hannah E. Smithson, Manuel Spitschan

## Abstract

Light in nature is complex and dynamic, and varies along spectrum, space, direction, and time. While both spectrally resolved measurements and spatially resolved measurements are widely available, spectrally and spatially resolved measurements are technologically more challenging. Here, we present a portable imaging system using off-the-shelf components to capture the full spherical light environment in a spectrally and spatially resolved fashion. The method relies on imaging the 4*π*-steradian light field reflected from a mirrored chrome sphere using a commercial hyperspectral camera (400-1000 nm) from multiple directions and an image-processing pipeline for extraction of the mirror sphere, removal of saturated pixels, correction of specular reflectance of the sphere, promotion to a high dynamic range, correction of misalignment of images, correction of intensity compression, erasure of the imaging system, unwrapping of the spherical images, filling-in blank regions, and stitching images collected from different angles. We applied our method to Wytham Woods, an ancient semi-natural woodland near Oxford, UK. We acquired a total of 168 images in two sites with low and high abundance of ash, leading to differences in canopy, leading to a total 14 hyperspectral light probes. Our image-processing pipeline corrected small (*<*3 ^*°*^) field-based misalignment adequately. Our novel hyperspectral imaging method is adapted for field conditions and opens up novel opportunities for capturing the complex and dynamic nature of the light environment.

## 1. INTRODUCTION

### 1.1 Background

Light plays a critical role in both visual and non-visual physiological processes of nearly all organisms. To fully understand the adaptation of an organism’s photoreceptive system to its surroundings it is necessary to place it within the context of its light environment. The light sensed by an organism is determined by three main factors: the characteristics of the illumination source, the spectral reflectance of objects in the local environment, and the spectral sensitivity of the organism itself. In the latter case, there is substantial interspecific variation in the repertoire of photopigments whose spectral sensitivities govern the wavelengths of light that are detectable by an organism.^1^

The light environment is complex and dynamic, and is defined along spectral, spatial, directional and temporal dimensions. The majority of light in the natural environment originates from the sun filtered through the atmosphere, with a very small portion in select environments arising from bioluminescence. Natural light varies in intensity across the 24-hour cycle due to solar elevation, ranging from 0.0001 lux on a clear starlit night to 100,000 lux on a sunny day, with greatest changes occurring during twilight.^2^ The changes at twilight occur concomitantly with predictable alterations to spectral composition, specifically a progressive enrichment of short-wavelength light (*<*500 nm) resulting from the Chappuis effect.^2–4^ Due to the tilt of the Earth’s axis, the length of the illumination period also alters with season, such that the northern hemisphere experiences longer days in the summer and shorter days in the winter.

Illumination that reaches the earth’s surface is subsequently modified by the differential transmission, absorbance and reflectance of nearby materials, which further shape the light environment. The local light environment can be characterised as the light that reaches a point in space, from all directions. Spectral measurement of local light environments is the focus of this paper. We envisage two primary applications of such data: Firstly to model the light sensed by different organisms at this location, and secondly to quantify the visibility of an object at this location that is visible only by virtue of the light it reflects. Both applications enable new tests of the visual ecosystem - the physical light environment as sensed by the organisms that interact with it.

Since the light environment is determined not only by illumination but by the spectral reflectance of objects in the environment, seasonal variation in the cover and colour of vegetation can have significant impacts on an organism’s visual ecosystem. However, this variation has not been fully characterised in a spatially and spectrally resolved fashion. Furthermore, it is not yet understood how this interacts with diurnal and seasonal changes in natural illumination, as well as the temporal and spatial niche occupied by an organism. Here we introduce a system for spectral measurement of local light environments, which is sufficiently portable for use in field research, and present a data sequence collected in deciduous woodland over a seven-week period in English springtime.

### 1.2 Methods for measuring directional lighting

Previous efforts to characterise directional light have used bespoke devices, such as the sphere-shaped multidirectional photometer that acquires intensity measurements from 64 angles.^5^ In principle, it is possible to add more photometers to increase the angular/directional resolution, but this introduces engineering complexity.

An alternative approach, popular in computer graphics, is to use a light probe that captures the light impinging on a single point in a scene.^6^ Light probes are used to generate environmental illuminant maps that allow rendering of light hitting a surface at the target location. There are two primary methods by which to capture a light probe - one is to use a camera with a so called ‘fish-eye’ lens, that maps wide angles to the sensor-plane of the camera, and the other is to photograph a mirror-sphere. Both methods require photographs taken from a small number of directions and the resultant images to be “unwrapped” and stitched to gain the full 4*π* steradian probe.^6^

### 1.3 Characterising natural light

Multiband or ‘multispectral’ imaging systems acquire spatially resolved images in multiple spectral channels, each sensitive to a different range of wavelengths. For example, dSLR cameras typically have three channels (RGB). The ability to recover spectra from multispectral images depends not only on the number of channels, but also the bandwidths of the channels. With low numbers of broadband overlapping channels, vastly different spectral distributions could produce very similar camera responses. Existing environmental light probes captured with standard dSLR cameras specify lights only in terms of RGB values, and not full spectral information. However, ecologically-relevant light probes should contain full spectral information over the range of wavelengths that are relevant to the biological process in question. Hyperspectral imaging systems sample light as a continuous function of wavelength. Hyperspectral cameras record spectra at each pixel, resulting in both high spatial and spectral resolution. Photographing mirror spheres with hyperspectral cameras has been demonstrated to accurately capture variations in directional lighting.^7^

## 2. METHODS

### 2.1 Approach

To characterise the hyperspectral light environment, we developed a portable imaging technique for capturing the 4*π* steradian light field in a spectrally and spatially resolved way. This imaging system was validated in Wytham Woods, a 426.5 ha area of ancient semi-natural woodland located 5 km outside Oxford (United Kingdom). Wytham Woods is a site of special scientific interest (SSSI) and has been the subject of extensive scientific research since it came under the ownership of the University of Oxford in 1942.^8^

We selected two sites within Wytham Woods for data collection, both of which were sampled on seven separate visits between April and June 2021. Both selected sites are part of an ongoing study exploring the impacts of the fungal pathogen *Hymenoscyphyus fraxineus* on European Ash (*Fraxinus excelsior*) mortality (“ash dieback”). One site has high ash density, whereas the second has been experimentally manipulated to simulate accelerated ash dieback, therefore representing the woodland a few decades after the ash has disappeared. Ash dieback results in the development of canopy gaps, with canopy cover often declining by 15% within a single season.^9^ This makes these sites interesting locations in which to validate our method since the data can be linked to changes in the light environment resulting from ash dieback, and the knock-on effects this will have on the structure and diversity of this ecosystem.

### 2.2 Hyperspectral Imaging System

The hyperspectral imaging system is shown in Figure 1a. It utilises a SpecimIQ push-broom hyperspectral imaging spectrometer (Specim; Oulu, Finland, 512 *×* 512 px resolution, 400-1000 nm, 204 spectral channels; 31 *×* 31 ^*°*^ FOV) to image a 3 in (76.2 mm) diameter chrome sphere, with an etched equator (AISI 52100 chrome steel ball, Simply Bearings Ltd, Lancashire, UK) that mirrors the surrounding environment. An imaging support system was developed to hold the hyperspectral camera, chrome sphere, and a white card. The system has two main components: a stationary stand (50 mm in height) and a rotating arm (800 mm in length). The chrome sphere was positioned on the stationary stand in the middle of the rotating arm. The hyperspectral camera sits at one end of the arm and images the chrome sphere against a white card, to simplify sphere extraction in post-processing (see Figure 1). Weights were used to counterbalance the camera, and the entire imaging system was mounted on a Manfrotto tripod.

**Figure 1.**
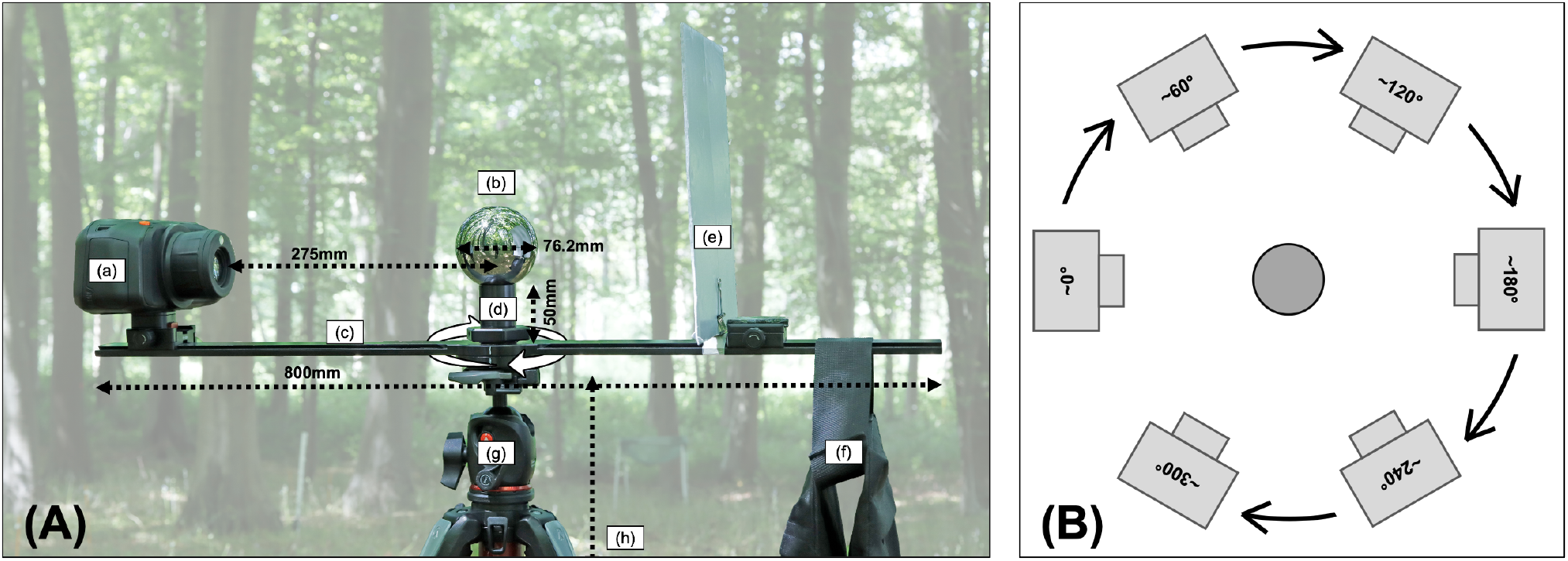
(A) Bespoke imaging system setup: (a) SpecimIQ hyperspectral camera, (b) 3 in (76.2 mm) chrome sphere, (c) Rotating arm, (d) Stationary stand, (e) White card, (f) Counterweight, (g) Tripod (h) Distance between ground and the tripod’s head, *∼*1360 mm. Distance between (a) and (b) is measured from lens to centre of chrome sphere. (B) Schematic illustration of measurement angles for one round of capture.

Our aim is to generate a full illumination map of the environment, measuring the light from every direction that falls at a particular point in the scene. The reflection of the environment in the sphere is largely compressed towards the edge of the sphere image so each acquisition predominantly captures the light reflected from one side of the sphere, thereby creating a blindspot behind the sphere. It is therefore necessary to image the sphere from multiple angles. The rotating arm enabled us to image the stationary sphere from six angles, separated by 60 ^*°*^ (see Fig. 1b), whilst maintaining a constant distance and elevation between the camera and the sphere. One round of acquisition comprised imaging the sphere from six azimuthal angles. This also allowed us to recover pixels lost from the reflection of the imaging system in the sphere, via the stitching together of these images in post-processing. The rotating arm increases the speed of data acquisition; minimising the change in the light environment across the sampling period.

### 2.3 Procedure

The imaging system was assembled in the field and the arm levelled manually with the help of an iPhone 12 mini and the app Angle Pro (available on the Apple App Store). To limit the loss of pixels, researchers stood behind the white card during data acquisition. As pixels containing the sky tend to be higher in intensity than ground pixels, we performed two rounds of data acquisition, prioritising ground and sky, respectively. This allowed us to visually assess and set an appropriate integration time that captured sufficient amounts of light whilst minimising under- or over-saturated pixels. On rare occasions, the longer integration times required by ground acquisitions were more susceptible to changes in light levels due to fluctuating cloud cover. These were accounted for by adjusting the integration time at different angles. A full imaging sequence for a given site took between 30 and 90 minutes to complete.

### 2.4 Image processing

#### 2.4.1 Absolute calibration

The SpecimIQ has a 204 (spectral) *×* 512 (spatial) push-broom sensor. Images with spatial resolution of 512 *×* 512 are created by shifting the angular orientation of the focal plane 511 times, to produce a 512 *×* 512 px *×* 204 band image. SpecimIQ performs calibrations by taking 100 images in an integrating sphere of known radiance and 100 dark images. The average dark pixel digital number (DN) is subtracted from each corresponding average light pixel DN. We presume but have not confirmed that Specim averages each spatial pixel that is shifted 511 times for each image. The calibration file is provided as a 1 *×* 512 *×* 204 band image. Once calibrated, the 204 values at each pixel in the calibrated image represent irradiance in µW ms cm^*−*2^ sr^*−*1^ nm^*−*1^.

To calibrate each image taken in the field, we subtract the dark reference DN image that is taken alongside each exposure from the raw DN exposed image. Once subtracted, we then multiply each pixel with the corresponding pixel from the calibration image. This results in an image with pixel values in µW ms cm^*−*2^ sr^*−*1^ nm^*−*1^.

#### 2.4.2 Pipeline to generate a hyperspectral illumination map

Figure 2 shows the image-processing pipeline. The goal of these processes was to create a full-field illumination map (light probe) from the 12 (6 angles *×* 2 integration times) individual hyperspectral component images

**Figure 2.**
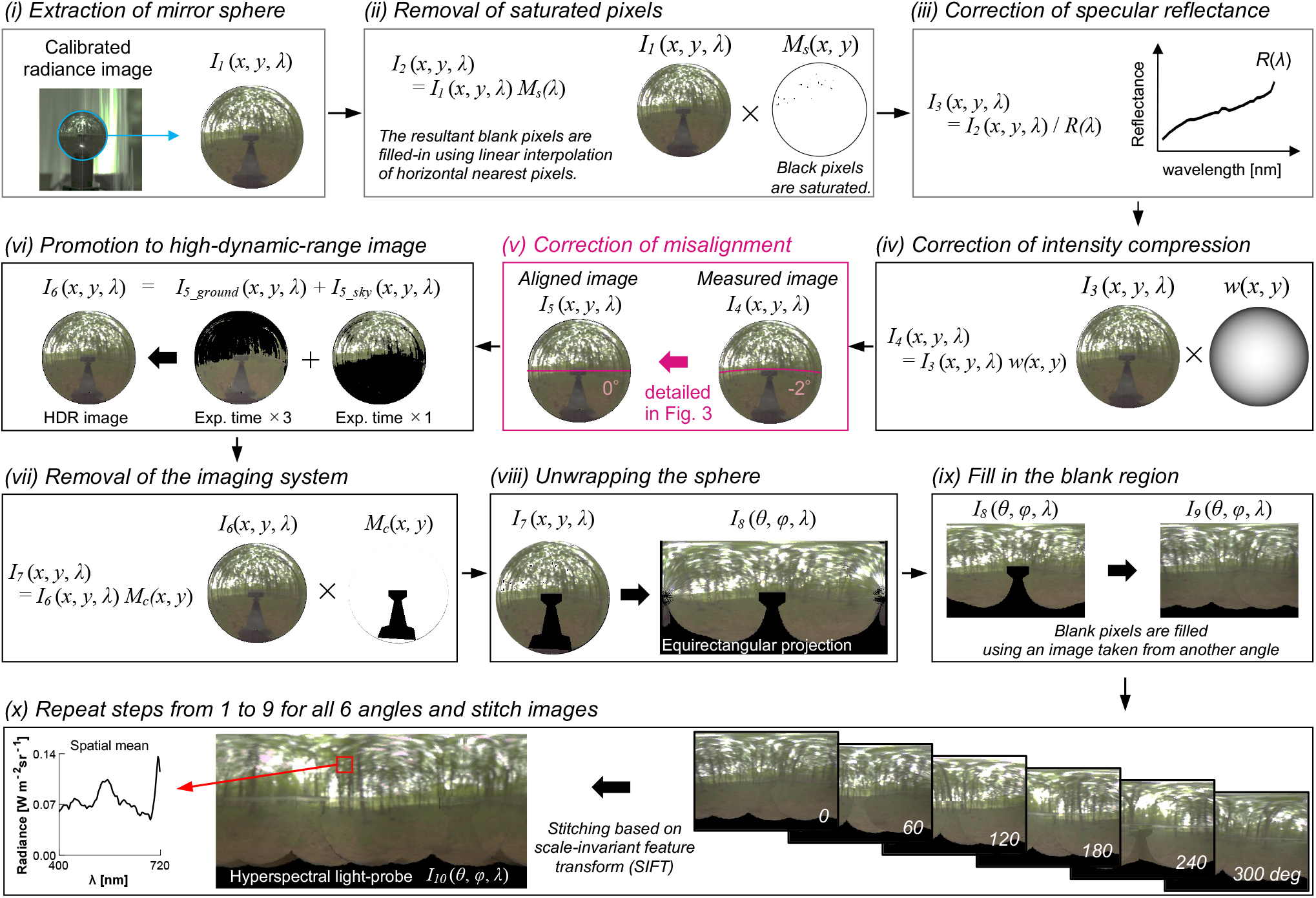
The image-processing pipeline to generate a hyperspectral light probe. (i) A sphere image was extracted from the raw image using a custom-built automatic sphere detection algorithm. (ii) Any saturated pixels were removed. (iii) Reflectance of the sphere was not flat over wavelength and thus it was corrected. (iv) We corrected the compression of intensity across the sphere. (v) We estimated the degree of misalignment, and applied the correction. (vi) Two images taken at different integration times were combined. (vii) The reflection of the imaging system was erased. (viii) The image was transformed from spherical coordinates to the equirectangular projection “unwrapping”. (ix) The blank region resulting from erasing the imaging system was filled with information from neighboring images. (x) Finally, images were stitched using an automatic feature-detection algorithm to create a full-panorama hyperspectral environmental illumination map.

First (Fig. 2i), we cropped out the sphere region *I* _1_(*x, y, λ*) from the calibrated radiance image.

Second (Fig. 2ii), using Eq. 1 we masked out saturated pixels using a binary mask *M* _*s*_(*x, y*), with which have zero or one at every wavelength corresponding to saturated and non-saturated pixels, respectively. After masking, the resultant blank pixels were filled-in by linear interpolation of horizontal nearest pixels.

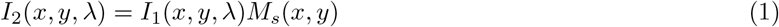

Third (Fig. 2iii), we corrected the spectrally non-flat reflection function of the mirror sphere *R*(*λ*) which was measured by spectrophotometer (UV-3600 Plus UV-VIS-NIR, Shimadzu, Kyoto, Japan^7^) based on Eq. 2.

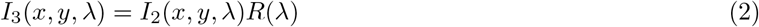

Fourth (Fig. 2iv), intensity compression in the image of the sphere was corrected by an empirically determined *w* (*x,y*) (Eq. 3).

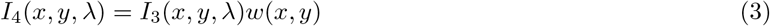

Fifth (Fig. 2v), although we used a rotating arm to minimise the misalignment of the imaging system, perfect alignment was challenging in a forest environment. Thus, we estimated the degree of misalignment and applied a post-hoc correction of the geometrical distortion introduced to the image (detailed in the next subsection). This ensured that the horizontal equator is mapped to the horizontal plane in the image.

Sixth (Fig. 2vi), we combined two images taken at the same angle but different exposure times to produce an image with higher dynamic range. Specifically we first converted the short-exposure image to luminance image and then masked out dim pixels in the image that had luminance values lower than 2 % of the highest luminance pixel. Then we filled in those regions with pixels of the ground image taken at approximately three times longer exposure time.

Seventh (Fig. 2vii), we masked out the camera region using a binary mask *M* _*c*_(*x,y*) based on Eq 4.

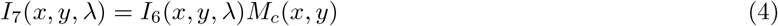

Eighth (Fig. 2viii), we converted the sphere image to equirectangular coordinates^10^ (transformation from a globe to a typical world-map), in which the horizontal and vertical coordinates represent the azimuth and the elevation angles, respectively.

Ninth (Fig. 2ix) we filled the blank regions introduced by erasing the imaging system taking the corresponding region from the image taken at a neighbouring measurement angle.

Finally (Fig.2x), we applied steps from (i) to (ix) to all other angles and stitched the processed images based on an automatic feature detection algorithm, the scale-invariant feature transform (SIFT^11^), to produce a hyperspectral illumination map. The extraction of image features was performed on the luminance-channel image, and we applied the identical stitching transformation to all wavelengths.

### 2.5 Estimation and correction of the misalignment of imaging system

For each measured sphere image, we estimated the misalignment of the imaging system relative to the target elevation of 0^*°*^, and applied a geometric correction. Our approach is to simulate the spatial distortion introduced to the sphere image due to camera misalignment using computer graphics techniques and to find a transformation that undoes the distortion. Figure 3 shows the procedure. First, using the rendering software Mitsuba,^12^ as shown in panel (a) we first generated 81 images of a mirror sphere with an etched black equator from camera positions ranging from *−*20 ^*°*^ to +20 ^*°*^ in 0.5 ^*°*^ steps. The image generated from 0 ^*°*^ corresponds to the ground-truth image. The mirror sphere was placed under a colorful light probe that varied in hue and saturation horizontally and vertically, respectively. Thus the mirror sphere rendered under this light probe exhibited distinct RGB values at each pixel, allowing us to find the transformation that needs to be applied to misaligned images. Specifically, as shown in panel (b), we created a displacement map for each of 80 incorrect angles, by finding for each pixel in the incorrect image (e.g. *−*20 ^*°*^) the pixel in ground-truth image (i.e. 0 ^*°*^) with the closest color. Then, as shown in panel (c), for a given measured sphere image, we estimated the degree of camera misalignment from projection of the equator in the measured sphere image. Then, we referred to the displacement map made in panel (b) as a look-up table to undo the spatial distortion, resulting in the aligned image where the horizontal equator becomes a straight line.

**Figure 3.**
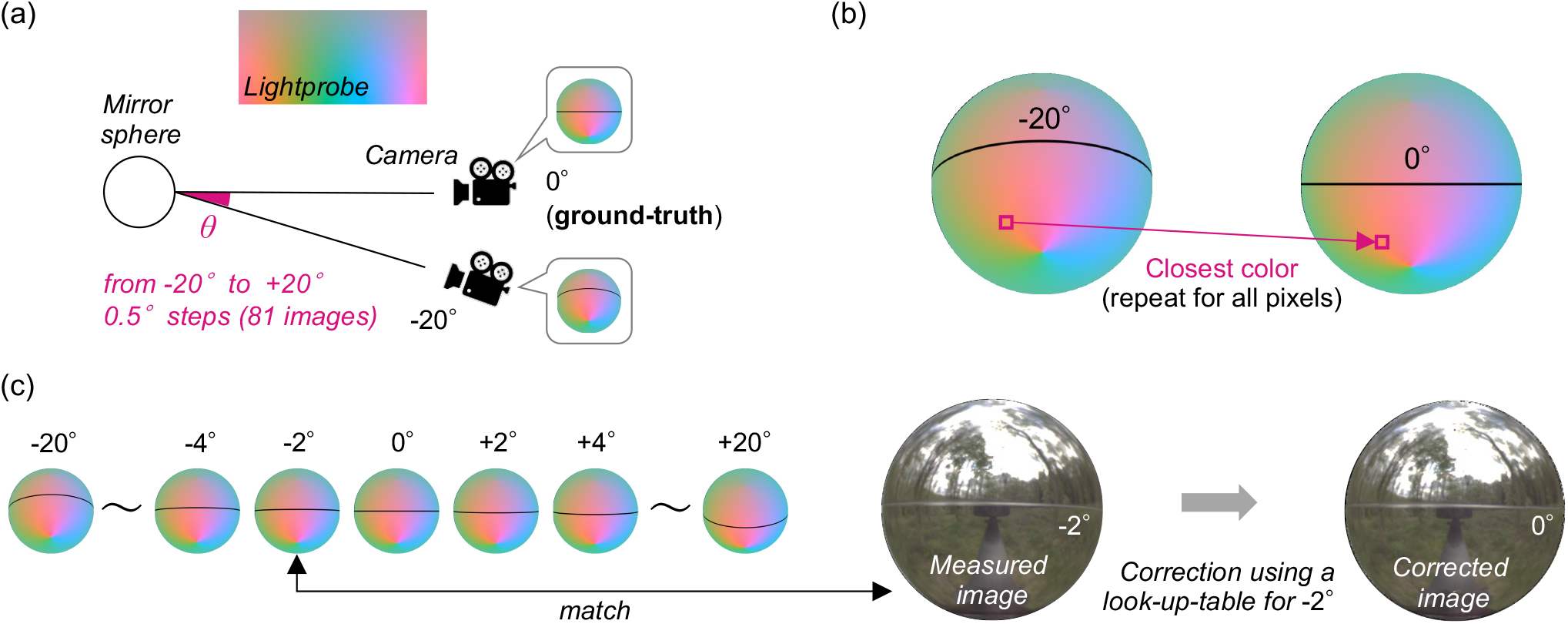
Estimation and correction of camera misalignment in elevation. (a) Generation of mirror sphere images from different camera elevations under a colorful light probe that differs in hue and saturation along azimuth and elevation directions (horizontally and vertically). (b) Each pixel in a sphere image stores a distinct RGB value, allowing us to map corresponding environmental features across images from different camera elevations. (c) Estimation of camera elevation using the distortion of the project of the etched equator, and correction using the corresponding look-up-table.

## 3. RESULTS

### 3.1 Captured images

In total we collected 168 images (14 environment maps *×* 6 angles *×* 2 integration times). Figure 4 presents an overview of hyperspectral images measured in this study. The lower part shows the 84 sphere images taken with shorter integration times aiming to capture the bright sky region. The upper pseudo-coloured spheres indicate intensity images at a range of wavelengths. Each sphere image was processed using the pipeline described in Section 2.4 to generate a full illumination map.

**Figure 4.**
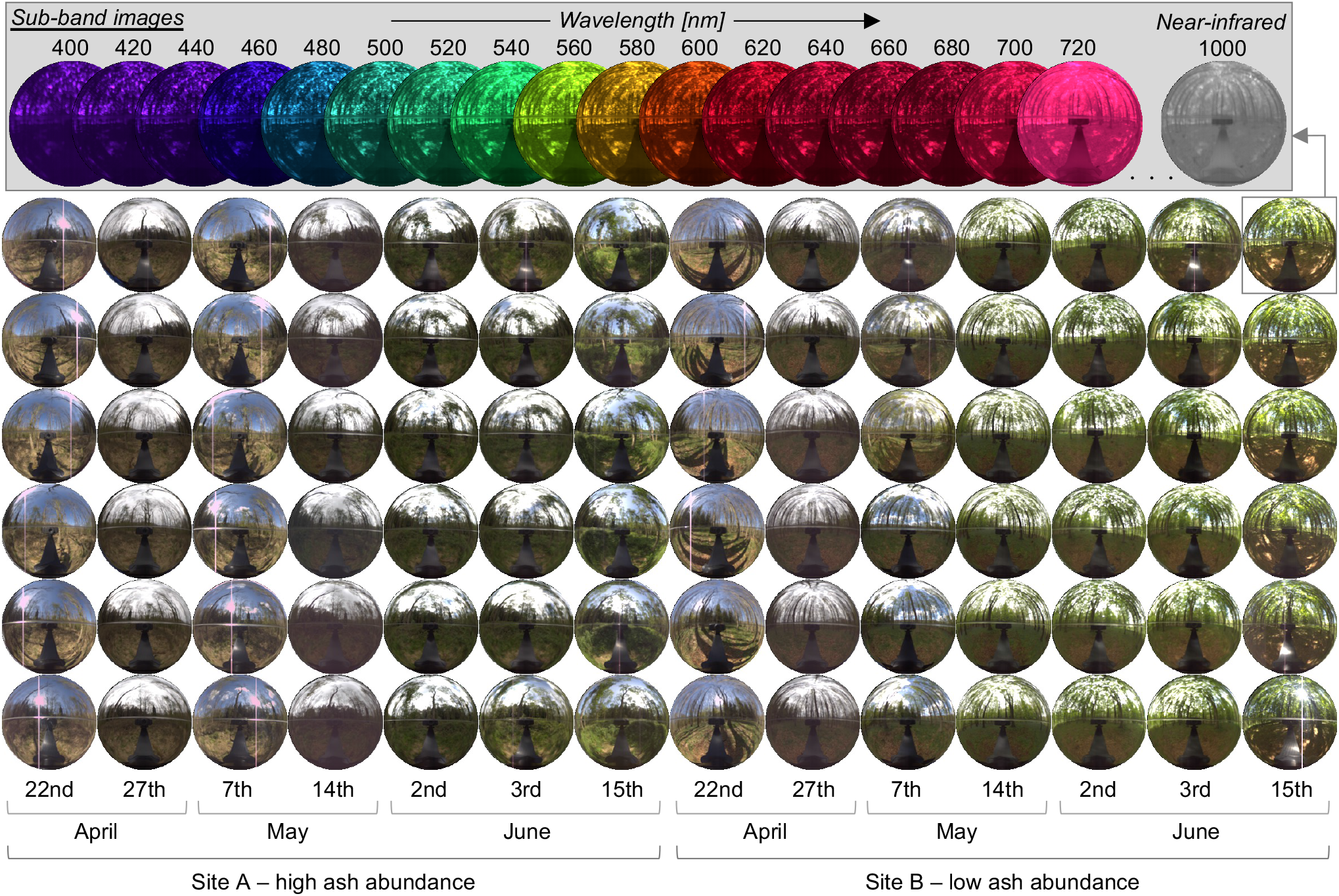
Collected hyperspectral sphere images in this study. The 84 circular images shows the sRGB representation of each acquisition and upper panel indicates wavelength sub-band images. The measurements were performed on the dates indicated at the bottom of the image. Only the 84 images taken with a shorter integration time are shown here, a further 84 images were collected with longer integration times.

### 3.2 Field-based misalignment

We evaluated the misalignment for all collected sphere images. We found that the majority of images were taken within the 1 ^*°*^ misalignment and the absolute error was always *<*3 ^*°*^ and always negative (i.e. the camera was lower than the sphere) as shown in Figure 5.

**Figure 5.**
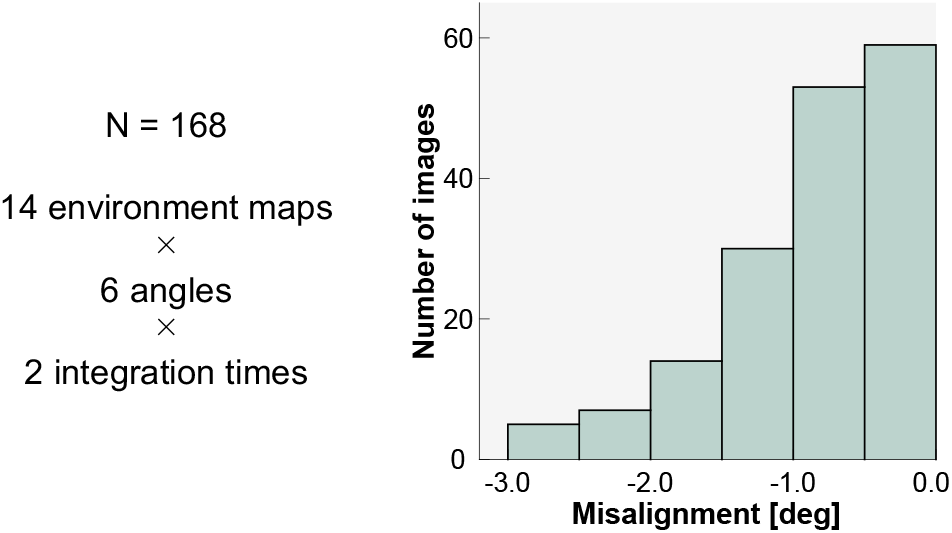
Histogram showing the observed camera elevations for the sample of 168 captured images.

**Figure 6.**
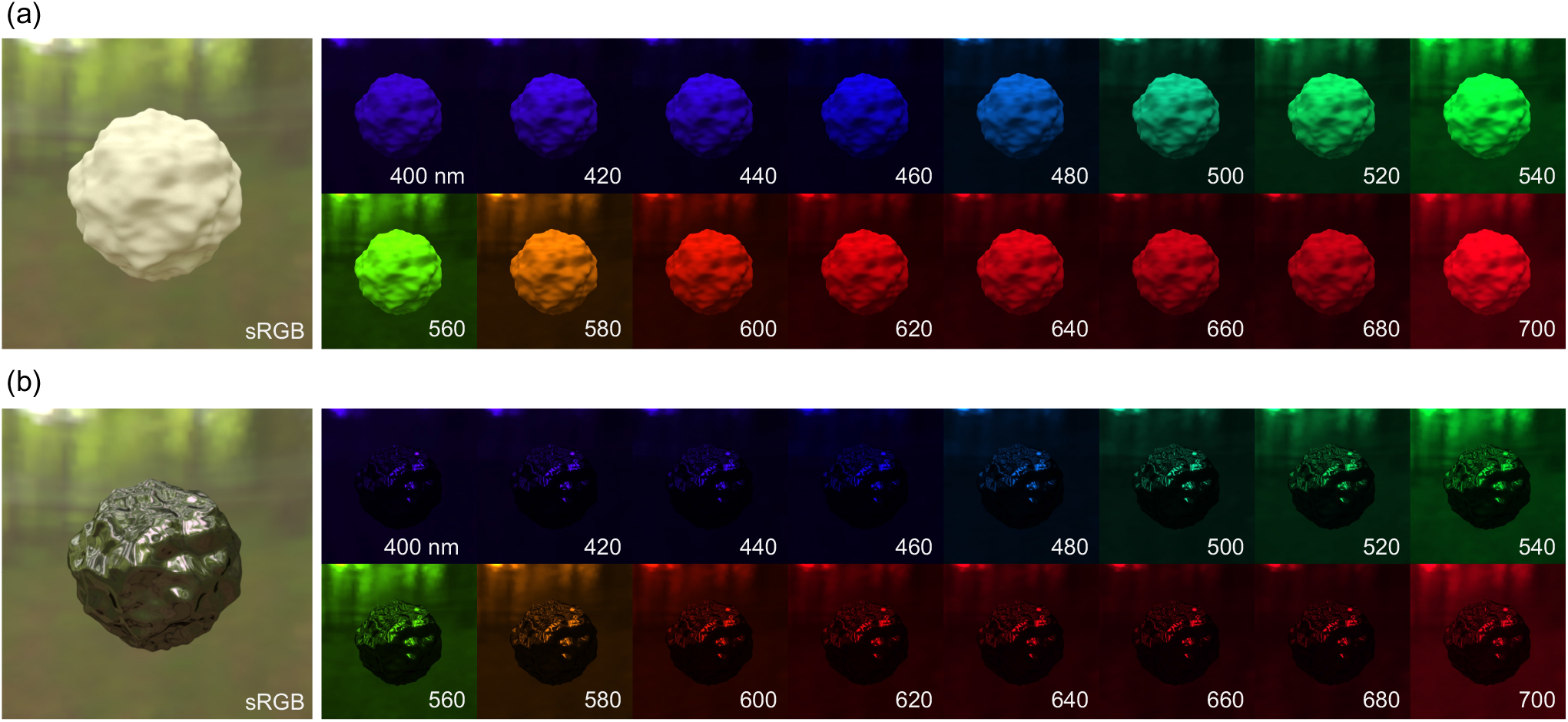
sRGB images and sub-band images from 400 nm to 700 nm with 10 nm steps of rendered objects under a processed hyperspectral light-probe. (a) A matte object with a diffuse reflection and (b) a mirror object that exhibits only a specular reflection.

### 3.3 Rendering using a hyperspectral light probe

To show an example application of the generated hyperspectral light probe, we used computer graphics techniques^6^ based on the rendering software Mitsuba^12^ to generate images of matte and glossy objects embedded in the environment. The synthetic objects have realistic appearance, and the spatial spectrum variation over the surface could be useful information to understand the daily visual inputs to organisms living in the environment, providing useful resources to researchers in different fields, from visual and circadian science to environmental science, who have interest in understanding the spectral nature of natural lighting environments.

## 4. LIMITATIONS

Our imaging system along with the image-processing pipeline is well-suited for capturing natural illumination in spatially and spectrally defined way. We have identified the following limitations of our method, which will be addressed in future work. First, the commercial, off-the-shelf camera we used only had a resolution of 512 *×* 512 px, which limits our ability to image fine spatial details in the environment. Second, due to the fixed-lens nature of the camera, the distance between camera and sphere was fixed, leading to a larger number of pixels representing our imaging system which had to be erased from the single images. Third, due to the sensitivity of the camera, sampling from one site took up to 90 minutes to complete, during which natural illumination could change due to, e.g., cloud cover. Fourth, due to weather conditions, our sampling of natural illumination could be biased due good-weather days. Finally, the wavelength range of the camera includes only visible and infrared wavelengths (400-1000 nm), which limits our ability to investigate the natural light environment as seen by organisms to those that have photoreception in that range.

## 5. CONCLUSION

Here, we aimed to develop a method to measure natural illumination in its dynamic and complex nature. We demonstrated the utility of a custom hyperspectral imaging system for the capture of environmental illumination in a circumscribed woodland environment at Wytham Woods. The imaging system was assembled from off-the-shelf imaging components, making it readily accessible for others, and integrated for effective and portable use in the field. Our custom post-processing pipeline addressed a range of issues arising from the measurement in a challenging field environment, including the misalignment of images taking from different angles, as well as the high-dynamic-range nature of natural scenes.

Our method is suitable to measure the light environment in complex light environments in a spectrally, spatially, directionally, and temporally defined way, and opens up novel opportunities for capturing light in nature. Capturing the light environment will enable the characterisation of visual and non-visual signals available to a wide variety of organisms.

## ACKNOWLEDGMENTS

This work was supported by an award from the John Fell Fund, University of Oxford (0008092). Laura Steel was supported by the BBSRC (Oxford Interdisciplinary Bioscience DTP; BB/S015817/1). Cecilia Dahlsjö was supported by NERC (NE/T007648/1). Alexander Shenkin was supported by NERC (NE/P012337/1). Takuma Morimoto was supported by the Wellcome Trust (218657/Z/19/Z) and Pembroke College, University of Oxford (Junior Research Fellowship). Manuel Spitschan was supported by the Wellcome Trust (204686/Z/16/Z).

This research was funded in whole or in part by the Wellcome Trust (204686/Z/16/Z, 218657/Z/19/Z). For the purpose of Open Access, the author has applied a CC BY public copyright licence to any Author Accepted Manuscript (AAM) version arising from this submission.

We thank Prof. Tom McLeish for discussions of the correction of the intensity compression imposed by the spherical imaging geometry, Prof. Nick Holliman for discussions on correcting image misalignment, and Mike Tacon from the Department of Physics workshop for manufacturing the rotating arm.

## Notes

### Competing Interest Statement

The authors have declared no competing interest.

